# Stability of Neural Oscillations Supports Auditory-Motor Synchronization

**DOI:** 10.1101/2023.10.19.563062

**Authors:** Rebecca Scheurich, Valentin Bégel, Ella Sahlas, Caroline Palmer

**Affiliations:** Department of Psychology, McGill University, 1205 Dr Penfield Ave, Montreal, QC, Canada H3A 1B1; Now at Institut des Sciences du Sport Santé de Paris (I3SP), Paris Cité University, 1 rue Lacretelle, 75015, Paris, France

**Keywords:** auditory perception, auditory-motor integration, synchronization, neural dynamics, recurrence quantification

## Abstract

Previous findings suggest that musical training leads to increased coactivation of auditory and motor brain networks, as well as enhanced auditory-motor synchronization. Less is known about the temporal dynamics of auditory-motor network interactions and how these temporal dynamics are shaped by musical training. The current study applied Recurrence Quantification Analysis, a nonlinear technique for characterizing the temporal dynamics of complex systems, to participants’ neurophysiological activity recorded via electroencephalography (EEG) during an auditory-motor synchronization task. We investigated changes in neural predictability and stability with musical training, and how these changes were related to synchronization accuracy and consistency. EEG was recorded while musicians and nonmusicians first tapped a familiar melody at a comfortable rate, called Spontaneous Production Rate (SPR). Then participants synchronized their taps with an auditory metronome presented at each participant’s SPR and at rates 15% and 30% slower than their SPR. EEG-based outcomes of determinism (predictability) and meanline (stability) were compared with behavioral synchronization measures. Musicians synchronized more consistently overall than nonmusicians. Both groups of participants showed decreased synchronization accuracy at slower rates, and higher EEG-based determinism (predictability) at slower rates. Furthermore, neural meanline (stability) measures correlated with synchronization consistency across all participants and stimulus rates; as neural stability increased, so did synchronization consistency. Neural stability may be a general mechanism supporting the maintenance of synchronization across rates, which may improve with musical training.

## Introduction

Musical behaviors across a wide range of contexts, from synchronizing body movements with a musical beat to adapting one’s performance tempo in an ensemble, require fine temporal coordination of auditory and motor systems. Given the auditory-motor coupling necessary to successfully carry out musical behaviors, there has been much interest in the relationship between musical experience and auditory-motor synchronization abilities. More hours of weekly music practice are associated with greater synchronization accuracy for highly demanding complex rhythms [1]. More years of musical training are related to increased auditory-motor flexibility: Musicians synchronize more accurately and adapt more efficiently than nonmusicians across a range of rates [2,3]. Specific aspects of musical experience may also play an important role in shaping auditory-motor synchronization abilities. Individuals with specialized percussion training show superior synchronization compared to their non-percussionist musically trained counterparts (e.g, [4,5]). Despite these findings supporting changes in auditory-motor processes with musical training, the neural mechanisms underlying these changes are still not fully understood. The current study examines the neural dynamics underlying enhanced auditory-motor synchronization and the impact of musical training on these dynamics.

There has long been an interest in the structural and functional changes in the brain that accompany musical training [6]. This research has pointed to specific changes in auditory and motor cortices as well as their interactions. Structurally, musicians show increased volume of the auditory cortex, which is related to greater functional connectivity at rest between auditory and motor regions [7]. Musicians who begin training earlier in life also show greater gray matter volume in the ventral premotor cortex [8], a region that appears to play a critical role in auditory-motor integration [6, 9]. This increased gray matter volume has been correlated with increased synchronization accuracy [8].

Additional studies suggest that increased integration between auditory and motor networks can occur with short-term musical training that may support enhanced auditory-motor synchronization. Using fMRI, Lahav et al. [10] examined changes in auditory-motor networks of nonmusicians after individuals were trained to play a melody on a keyboard by ear. Following training, listening to the melody (in the absence of movement) elicited activity in motor regions. Similar work with EEG has also shown that, following a period of musical training, nonmusicians had greater auditory motor coactivation in response to separate auditory and motor tasks [11]. This result is consistent with EEG source localization findings with musicians: Pianists show greatest activity in motor cortices in response to hearing auditory sequences that they had previously produced, relative to melodies they had previously heard [12].

Despite substantial evidence to suggest that musical training increases integration of auditory-motor networks, little research has examined the temporal dynamics of changes in neural network activity. One study classified listeners’ EEG patterns for auditory rhythms using convolutional neural networks [13], a deep learning technique that assumes stationarity of the data (an unchanging mean and variance over time). Nonlinear analysis techniques, well-suited for periodic (nonlinear) neural measures, provide quantification of the temporal dynamics of neural activity without an assumption of stationarity. Recurrence Quantification Analysis (RQA) is a promising avenue for examining nonlinear regularities in complex neural systems (e.g., [14]). RQA identifies recurrences in patterns of neural activity over time without making assumptions about data stationarity, and thus is an ideal candidate method for exploring the temporal dynamics of auditory-motor network integration.

Most applications of RQA to EEG signals can be found in clinical research, such as classifying epileptic patients’ EEG activity [15] or evaluating patients’ levels of consciousness [16]. Only a few studies have applied RQA to investigate auditory-motor behaviors and neural activity in musicians and nonmusicians. Comparisons of auditory-motor synchronization in musicians and nonmusicians using RQA have shown differences in the predictability and stability of their synchronization behaviors [2]. Additionally, RQA applied to intra- and inter- brain EEG dynamics during a joint rhythmic synchronization task suggests that temporal dynamics are shaped by the partners’ rhythmic interactions [17]. Thus, we expect that RQA can characterize auditory-motor network interactions in musicians and nonmusicians, and how these interactions support synchronization.

The current study investigated effects of musical training and stimulus rate on the temporal dynamics of neural networks in auditory-motor synchronization. Due to its high temporal resolution suitable for the study of temporal dynamics, electroencephalography (EEG) was used to measure cortical activity as musicians and nonmusicians completed auditory-motor synchronization tasks. To calibrate synchronization rates to each individual, participants first produced a familiar melody at a comfortable rate (called Spontaneous Production Rate) in the absence of external rate cues, a task that provides measures of an individual’s optimal temporal stability [2,18]. Participants then synchronized their tapping of the melody with a metronome set to their SPR and to slower rates. Slower (as opposed to faster) rates were chosen based on findings that musical training provides larger synchronization advantages at slower rates [2, 19].

Based on previous findings, we predicted that musicians would show greater synchronization accuracy and consistency than nonmusicians, and that synchronization accuracy and consistency would decrease for all participants at slower rates [2,18]. We examined neural dynamics during the synchronization tasks by applying auto-Recurrence Quantification Analysis of EEG activity at electrodes previously shown to elicit highest power in auditory-motor synchronization tasks [17, 20]. We predicted that musicians would show greater neural predictability (determinism) and stability (meanline) across synchronization rates than nonmusicians. If neural stability during synchronization tasks supports behavioral synchrony, then greater neural stability should be related to greater synchronization consistency, a behavioral measure of stability.

## Materials and methods

### Participants

Twenty-four participants provided informed consent to participate in the study. Twelve musically trained individuals (mean age = 25.09 years old, SD = 3.81; 6 female, 3 male, 2 other, 1 undisclosed) and 12 untrained individuals (mean age = 23.17 years old, SD = 3.86; 8 female, 3 male, 1 undisclosed) participated in the experiment. The participant groups did not differ in age, *t* (21) = 1.20, *p* = 0.24. Percussionists were excluded from the current study based on previous evidence suggesting differences between percussionists and non-percussionists in auditory-motor tasks (e.g., [21]). Musicians had at least 6 years of private instrumental instruction (mean = 11.33 years, SD = 3.06). Nonmusicians had less than 2 years of private music instruction, none of which had occurred in the past 6 years (mean = 0.06 years, SD = 0.15). All participants were right-handed as determined by the Edinburgh Handedness Inventory [22], had normal hearing in the frequency range of experimental stimuli as determined by an audiometry screening (< 30 dB HL threshold for 250-1000 Hz), reported no neurological disorders, and were not taking medication affecting the central nervous system at the time of participation. Participants were recruited during 02/20/2020 to 07/14/2022 (with COVID-related interruptions) and participants’ written consent was obtained.

### Stimulus Materials and Equipment

Participants’ hearing thresholds were determined using a Maico MA 40 audiometer. During the tapping tasks, participants tapped on the force-sensitive resistor (FSR) of an Arduino that controlled and delivered auditory feedback based on participants’ taps. Tones were delivered in a piano timbre from an Edirol Studio-Canvas SD-80 tone generator through EEG-compatible earphones (Etymotic ER-1, Etymotic Research, Inc.). The familiar stimulus melody produced by participants’ taps was “Twinkle, Twinkle Little Star,” composed in G Major. Auditory feedback from the metronome in the synchronization task was delivered in a woodblock timbre (pitch = E5). MIDI data were collected using FTAP on a Dell computer running Linux [23]. FTAP software was modified to integrate Lab Streaming Layer (LSL; [24]) so that triggers from FTAP were sent to a second Dell computer running LSL in Windows 10 over the local area network, as described and implemented previously [17,25].

### EEG Data Recording

EEG data were recorded at a 512 Hz sampling rate with a 64-channel BioSemi ActiveTwo system (BioSemi, Inc.). EEG was grounded using the common mode sense (CMS) and driven right leg (DRL) electrodes. Sixty-four scalp electrodes were placed according to the 10-20 system.

### Design

Each participant performed the spontaneous production rate and auditory-motor synchronization tasks in a fixed order so that the synchronization stimulus rates could be calibrated to each participant. The stimulus rates were set equal to the participant’s SPR (computed as the mean inter-onset interval (IOI) in ms), 15% Slower than their SPR, or 30% Slower than their SPR. The experiment had a mixed design with a between-subjects independent variable of Group (Musician and Nonmusician) and a within-subjects independent variable of Rate (SPR, 15% Slower, and 30% Slower). All participants completed the rate conditions in the same order, starting with the SPR followed by the 15% Slower and then 30% Slower rate conditions. This constant order was chosen to ease participants into the task, starting with their most comfortable rate.

### Procedure

After providing written informed consent upon arrival to the lab, participants underwent an audiometry screening. Participants who passed the audiometry screening and then reported familiarity with the experimental melody moved on to the remaining experimental tasks.

Participants were outfitted with the EEG cap and electrodes and seated in front of a fixation cross. The tapping sensor was placed on a table next to participants where they could rest their arm while tapping. Participants were then given earphones to complete the SPR task. Participants were instructed that they should tap the rhythm of “Twinkle, Twinkle Little Star” while focusing their gaze on the fixation cross in front of them. They were told that each time they tapped, a tone of the melody would sound and that they had control over when the next tone in the melody would sound, so they should tap the rhythm of the melody. Participants completed one practice trial in which they tapped the melody once through to ensure that they had understood the instructions. They then completed a second practice trial in which they tapped the melody 3.5 times through until they no longer heard the sound of their taps, indicating the end of the trial. Once participants were comfortable with the task, they were instructed that they should tap the rhythm of the melody at a comfortable and steady rate. Participants then completed three experimental trials in which they tapped the melody 3.5 times through until they no longer heard the sound of their taps, indicating the end of the trial. Following the SPR task, participants completed several questionnaires while their SPRs were computed for use in the synchronization task (see **Data Analysis**). These questionnaires assessed participants’ musical background and handedness.

Finally, participants completed the synchronization task. Participants were instructed that they would again be tapping the rhythm of “Twinkle, Twinkle Little Star” as they did before, but that this time they would hear a metronome and their goal was to synchronize their tapping with the metronome while focusing their gaze on the fixation cross in front of them. Participants first completed a practice trial in which they began synchronizing with the metronome after the first eight metronome clicks and continued to synchronize with the metronome for 3.5 repetitions of the melody until they no longer heard the sound of their taps, indicating the end of the trial.

Participants then completed three experimental trials. This procedure was repeated for all rate conditions. Participants performed additional perceptual (auditory) and motor tasks; for the purposes of this paper, only the auditory-motor synchronization task is included. Participants were debriefed following the synchronization task and given a small compensation for their time. The full study lasted approximately 2.5 hours.

### Behavioral Data Analysis

Participants’ SPRs were calculated as the mean IOI for the middle two repetitions across trials from the SPR task, to avoid tempo changes typically seen at melodic phrase beginnings and endings [26–28]. Half notes were interpolated and outlier IOIs more than three standard deviations away from the mean were removed (Musicians = 1.88% of total IOIs identified as outliers; Nonmusicians = 2.08%). The resulting SPRs were then used to determine the metronome rates for each participant in the synchronization task.

To examine synchronization performance in the synchronization task, tap and metronome onsets were first aligned in each trial using a nearest neighbor approach (e.g.,[29,30]). Signed asynchronies were then calculated as the tap onset time minus metronome onset time, with negative values indicating that a tap preceded the metronome and positive values indicating that a tap lagged the metronome. Signed asynchrony outliers more than three standard deviations from the mean signed asynchrony were removed (Musicians = 1.57% of total signed asynchronies identified as outliers; Nonmusicians = 2.55%). To assess whether participants were synchronized above chance in each trial, signed asynchronies were then converted to radians and the Rayleigh test for circular non-uniformity was run on the radian values per trial. A significant Rayleigh test indicates a significant mean direction (i.e., unimodal synchronization pattern), whereas a non-significant Rayleigh test indicates no significant mean direction (i.e., no synchronization or multimodal synchronization pattern [31]). Following previous work [29, 32–34], trials in which the Rayleigh test failed to reach significance were removed from all subsequent behavioral and EEG analyses (Musicians = 0.93% of total trials; Nonmusicians = 6.48% of total trials). An additional two trials for two musicians and one trial for one nonmusician were removed due to technical measurement difficulties. The mean signed asynchronies and standard deviations of the signed asynchronies were then computed for each synchronized trial.

### EEG Preprocessing and Data Analysis

EEG data were preprocessed using EEGLAB [35]. Data for all trials and tasks were first concatenated together and re-referenced to the common average across electrodes. Electrodes with poor signal quality were then detected by visually inspecting distributions of deviations from mean activity. Those with no deviation from mean activity or very large deviations from mean activity were identified as flat and noisy electrodes, respectively, and removed. The data were then filtered between 1 Hz and 40 Hz using a Hamming windowed sinc FIR filter (high and low pass filter order = 1000), segmented into 1-second epochs, and pruned for non-stereotypical artefacts. Data were then submitted to extended infomax Independent Component Analysis (ICA). Resulting ICA components reflecting eyeblink and lateral eye movement artefacts were visually identified from component topomaps and removed from the unfiltered data. Finally, electrodes removed due to poor signal quality were spherically interpolated.

EEG data were then prepared for auto-recurrence quantification analyses. Following previous research showing maximal power at C1 during an auditory-motor synchronization task [17], C1 was selected as input to auto-recurrence quantification analysis. Data recorded from C1 during the synchronization task were first filtered at the beat frequency corresponding to the metronome rate in each subject’s rate conditions. Filter cutoffs were defined as two standard deviations above and below the tapping frequency in the corresponding rate condition for each participant to account for timing deviations from the metronome beat frequency. A Hanning windowed sinc FIR filter was used (high and low pass filter order = 1000). A 14.5 second epoch, corresponding to the shortest melody repetition, was then extracted from the start of the second melody repetition for each significantly synchronized trial, to capture participants’ most stable performance. Data were finally *z*-scored per epoch.

Auto-recurrence analyses were conducted using the Cross Recurrence Plot Toolbox in MATLAB [36]. Parameters were optimized per rate condition and participant. The optimal delay parameter, determined using Average Mutual Information (AMI), was selected by identifying the first local minimum, or the first delay at which the least information is shared with the first point in the time series. The optimal number of embedding dimensions was determined using False Nearest Neighbours (FNN) by identifying the number of dimensions for which the number of false nearest neighbours in the phase space is minimized. The optimal number of embedding dimensions was determined to be the same across rate conditions and participants (selected embedding dimensions = 4), consistent with a previous study that implemented the same pipeline [17].

Rate manipulations can create a potential confound in the interpretation of observed differences in auto-recurrence outcomes. The same-sized time window applied to slower and faster rates may yield greater recurrence at the faster rate simply because there are more cycles of that rate that fall within the analysis window. To control for this potential confound, recurrence rate was fixed to 10% following previous studies [17, 36]. The Thieler window and the minimum diagonal and vertical line lengths were set to 23 samples for all rate conditions and participants, corresponding to 1/2 of the smallest optimal delay across rate conditions and participants (i.e., approximately 1/8 of the cycle of the fastest rate).

The auto-recurrence outcomes of interest were determinism and meanline. Sample recurrence plots are shown in Figure 1 alongside the time series EEG data, for one participant’s trial in the SPR rate condition (Figure 1a) and in the 30% Slower rate condition (Figure 1b) as a visual depiction. Determinism describes the proportion of recurrent points that fall along diagonal lines in the recurrence plot, and represents predictability in periodic systems [14]. Determinism is computed as shown in Eq. 1 [36], where *l* refers to the diagonal line length, *P*(*l*) refers to the histogram of the diagonal lines of length *l*, and *l_min_* refers to the minimum diagonal line length.

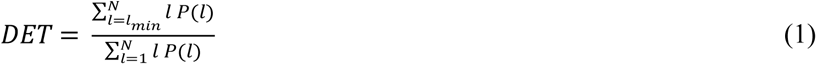

**Figure 1.**
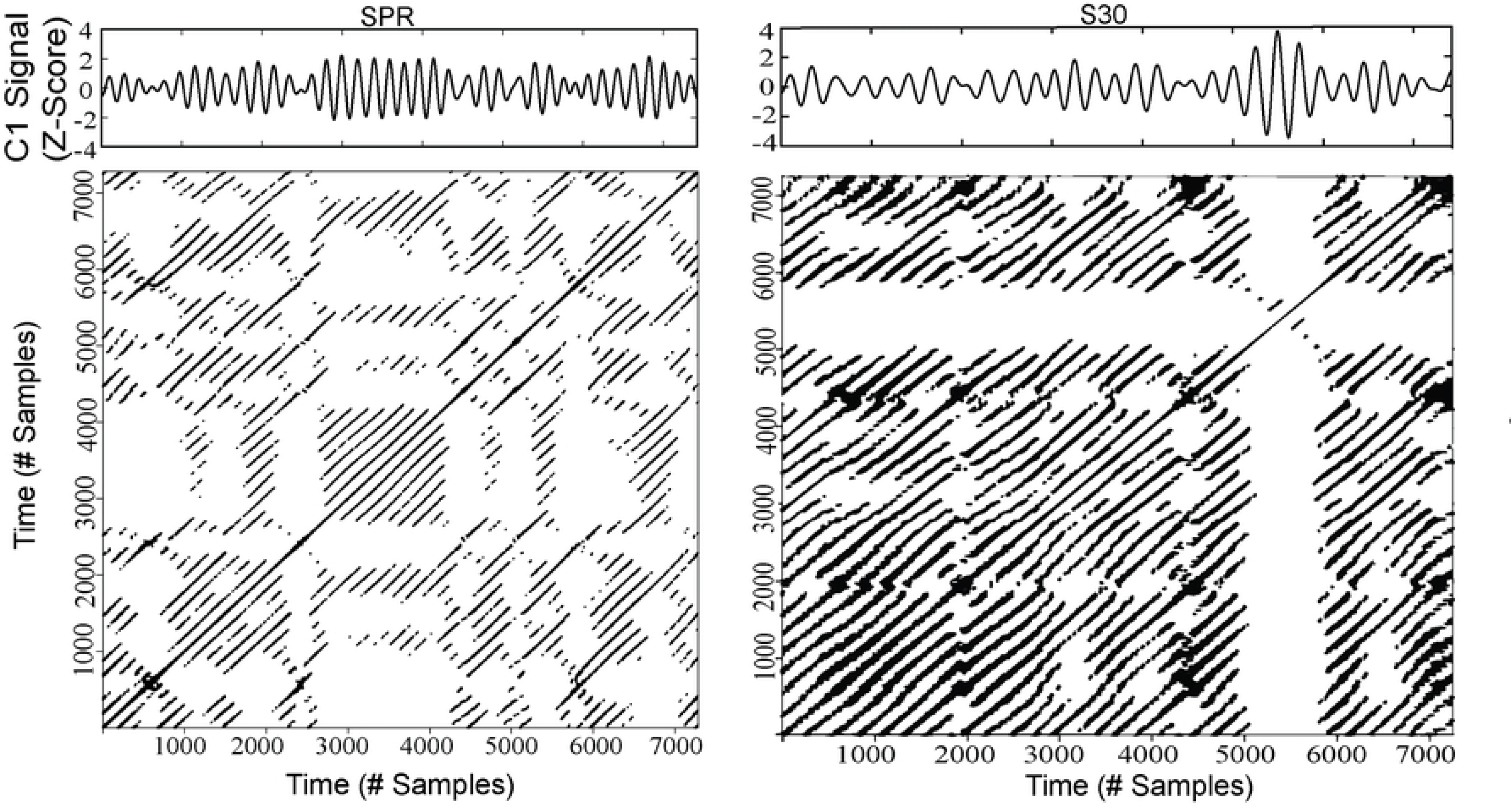
Sample auto-recurrence plots and corresponding time series from a musician’s EEG signal. Determinism is the proportion of recurrence points that fall on diagonal lines, and meanline is the average length of the diagonal lines. A: A single trial at the SPR rate. B: A single trial at the 30% Slower rate.

Meanline describes the average diagonal line length (# samples) in the recurrence plot and corresponds to the average length (or period) of repeating patterns [14, 37]. Meanline has been used as an index of a system’s stability [38–40] : longer lines (larger values) indicate greater stability. The computation of meanline is shown in Eq. 2 [36]. The main distinction between Eq. 1 and Eq. 2 is the denominator. The denominator in Eq. 2 represents the total number of diagonal lines, in contrast to Eq. 1 in which it represents the total number of recurrent points.

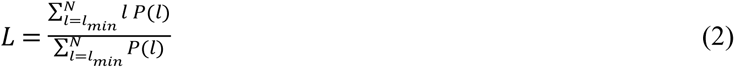

### Statistical Analysis

Analyses of variance (ANOVA) conducted by Rate and Group conditions are reported for both behavioral and neural measures, with effect sizes included. Posthoc comparisons are reported with Holm’s correction for number of tests. Robust ANOVAs are reported when tests of homogeneity of variance fail. Additionally, Pearson correlations are reported to examine brain-behaviour relationships between 1) synchronization stability (standard deviation of the signed asynchronies) and neural stability (meanline), and 2) synchronization accuracy (mean signed asynchrony) and neural predictability (determinism).

## Results

### Spontaneous Rates

Figure 2 shows participants’ mean SPR values, computed as mean intertap interval (ITI), ordered fastest to slowest (minimum = 302.95 ms ITI; maximum = 752.84 ms ITI). To examine whether SPRs varied across groups and trials, a two-way mixed ANOVA was run on the mean IOI with Group and Trial (1, 2, and 3) as factors. Results showed a significant main effect of Trial, *F* (2, 44) = 15.61, *p* < 0.001, η^2^ = 0.42. All participants sped up from trial 1 to trials 2 and 3 (Holm-adjusted *p’s* < 0.001 [41]), and from trial 2 to trial 3 (Holm-adjusted *p* < 0.05). Importantly, there was no significant main effect of Group, *F* (1, 22) = 1.17, *p* = 0.29, η^2^ = 0.05, and no significant interaction between Group and Trial, *F* (2, 44) = 0.15, *p* = 0.87, η^2^ = 0.01.

**Figure 2.**
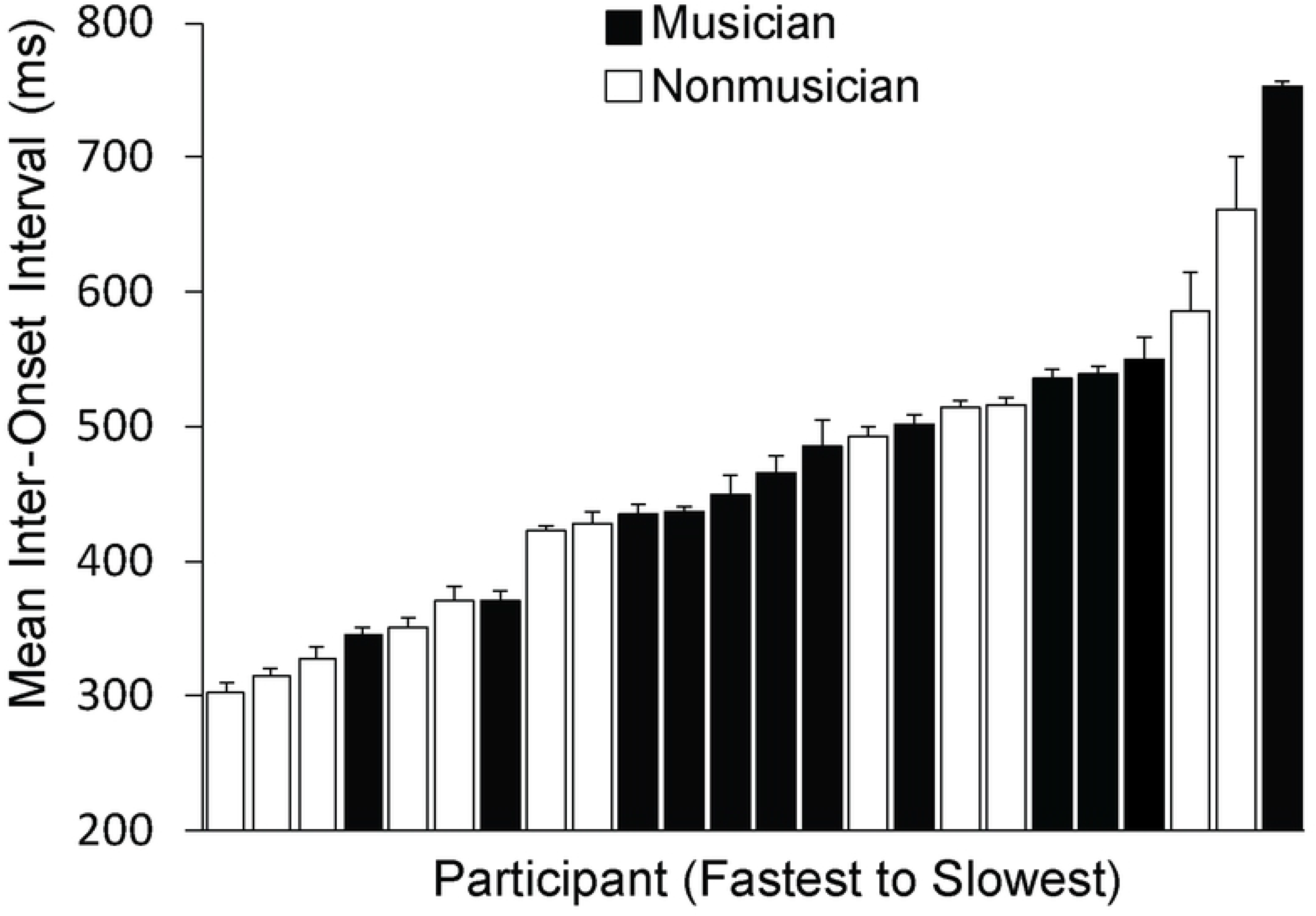
Distribution of participants’ SPR values (mean IOI, ms) ordered from fastest to slowest. Black bars represent musicians and white bars represent nonmusicians. Error bars represent standard error.

### Synchronization Accuracy

We next examined synchronization accuracy, measured via the mean signed asynchrony, by group and rate condition. As Levene’s [42] test indicated that homogeneity of variances was violated at the 15% Slower rate, *F* (1, 22) = 4.89, *p* < 0.05, a robust ANOVA using the trimmed means method [43] was run on the mean signed asynchrony with Group and Rate (SPR, 15% Slower, and 30% Slower) as factors. The significant main effect of Rate, *Q* = 6.32, *p* = 0.02, shown in Figure 3, indicated that signed asynchronies became more negative as the rate became slower than each participant’s spontaneous rate. There was no significant effect of Group, *Q* = 0.17, *p* = 0.68, and no significant interaction between Group and Rate, *Q* = 2.25, *p* = 0.16.

**Figure 3.**
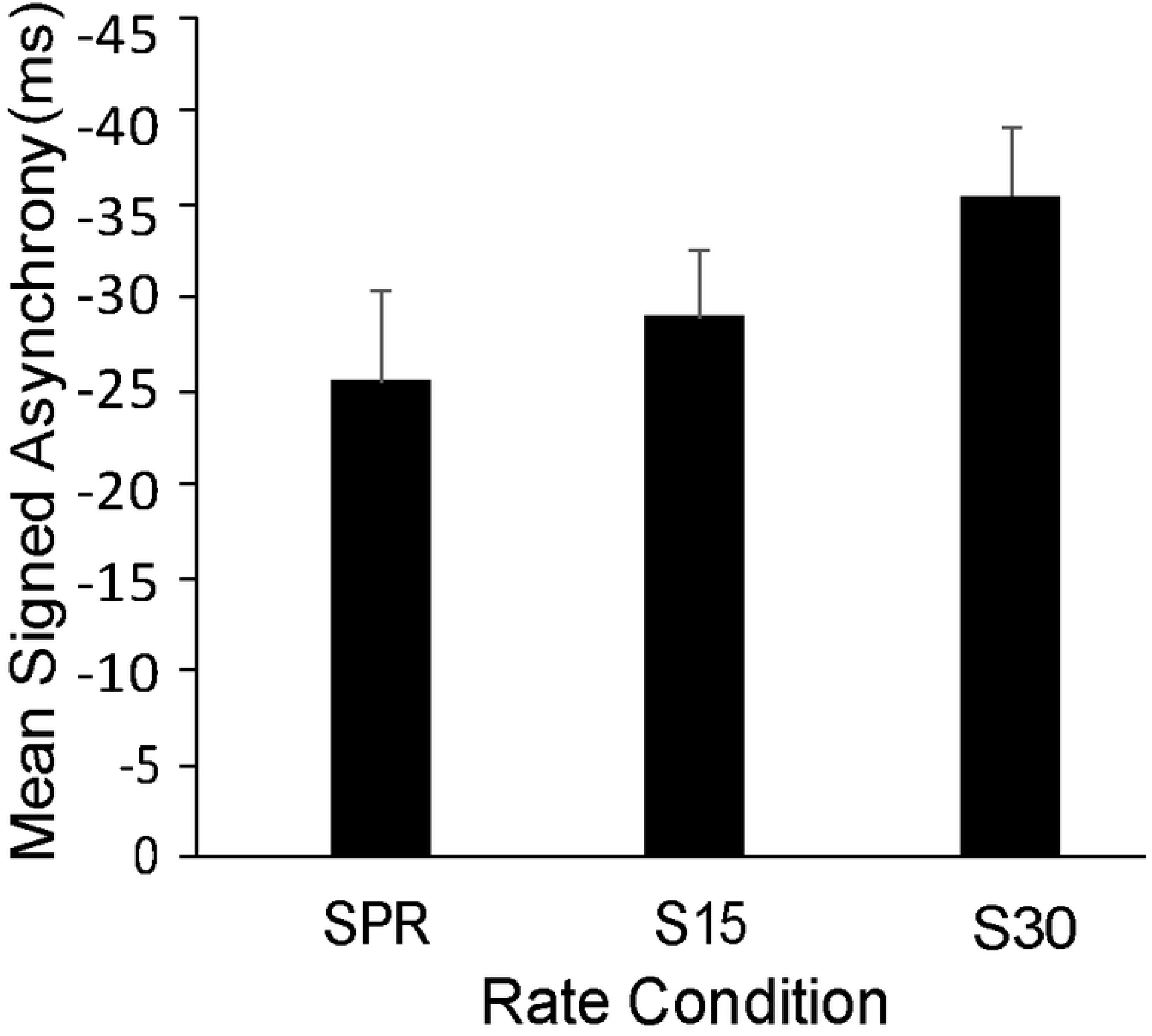
Mean signed asynchrony (ms) by rate condition. Error bars show standard error. *Synchronization Consistency.* We next examined synchronization consistency, measured via the standard deviation of the signed asynchronies, by group and rate condition. A two-way mixed ANOVA was run on the standard deviation of the signed asynchrony with Group and Rate as factors. Results showed a significant effect of Group, *F* (1, 22) = 15.90, *p* < 0.001, η^2^ = 0.42.

Musicians synchronized more consistently (smaller standard deviations; mean = 28.90 ms, SE = 2.99 ms) than Nonmusicians (mean = 59.90 ms, SE = 4.33 ms). There was no significant effect of Rate, *F* (2, 44) = 0.73, *p* = 0.49, η^2^ = 0.03, and no significant interaction between Group and Rate, *F* (2, 44) = 1.92, *p* = 0.16, η^2^ = 0.08.

### Auto-Recurrence Outcomes

We next assessed the predictability and stability of EEG activity during the synchronization task. We ran auto-recurrence quantification analysis on electrode C1 (representing auditory-motor network activity) and examined the outcomes of determinism (predictability) and meanline (stability). Two separate two-way mixed ANOVAs were run on determinism and meanline with Group and Rate as factors.

The first ANOVA on determinism showed a significant effect of Rate, *F* (2, 44) = 160.18, *p* < 0.001, η^2^ = 0.88. As shown in Figure 4, determinism was higher at the 15% and 30% Slower rates than at the SPR, and higher at the 30% Slower rate than at the 15% Slower rate (Holm-adjusted *p*’s < 0.001). There was no significant effect of Group, *F* (1, 22) = 1.36, *p* = 0.26, η^2^ = 0.06, and no significant interaction between Group and Rate, *F* (2, 44) = 0.85, *p* = 0.43, η^2^ = 0.04.

**Figure 4.**
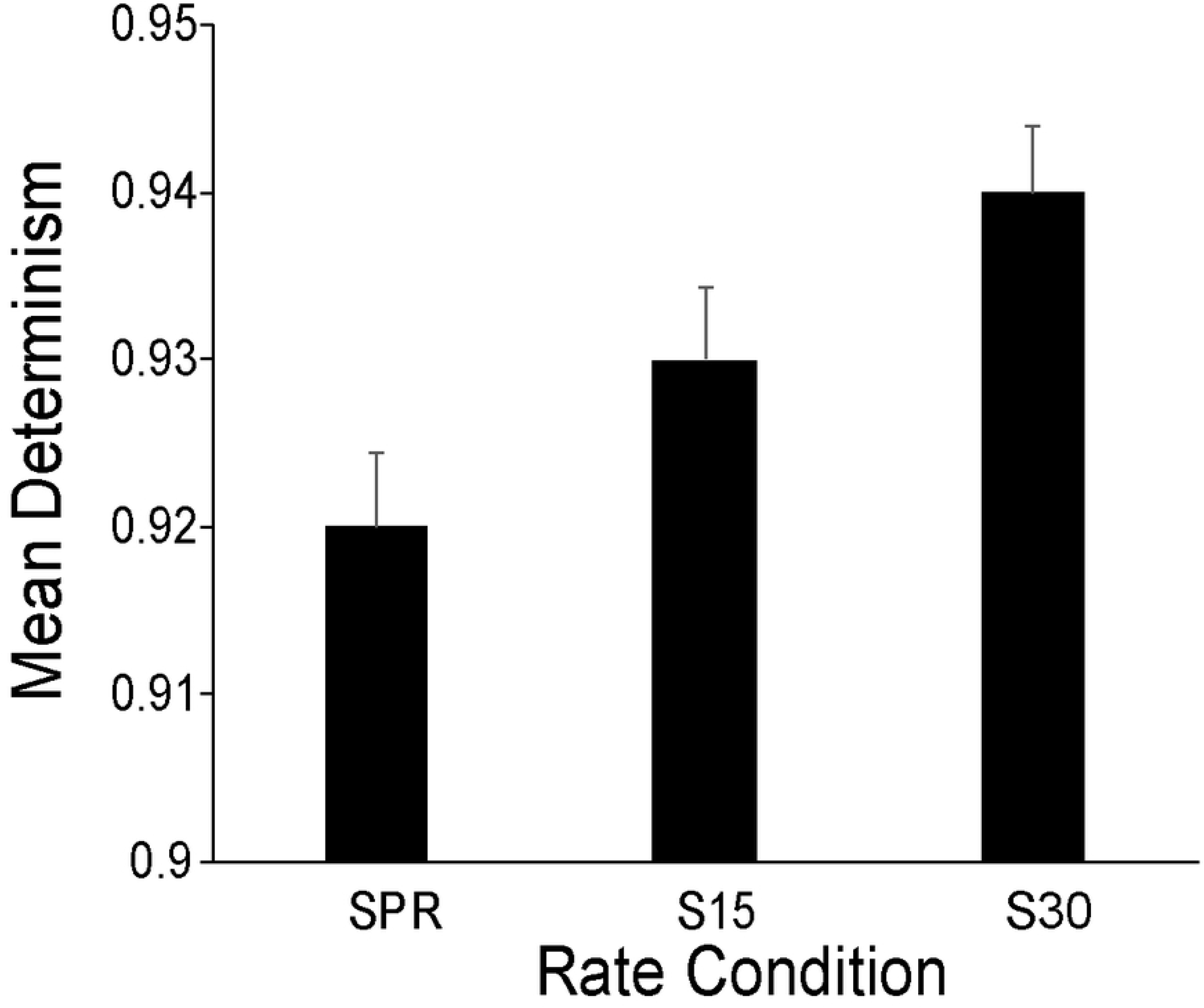
Mean determinism (predictability) in EEG signal by rate condition. Error bars show standard error. The second ANOVA on meanline showed that the effect of Group trended toward significance *F* (1, 22) = 3.71, *p* = 0.07, η^2^ = 0.14. Musicians tended to show greater neural stability (longer meanline; mean = 145 samples, SE = 1.82 samples) than Nonmusicians (mean = 139 samples, SE = 1.85 samples). There was no significant effect of Rate, *F*(2, 44) = 2.20, *p* = 0.12, η^2^ = 0.09, and no significant interaction between Group and Rate, *F*(2, 44) = 0.35, *p* = 0.71, η^2^ = 0.02.

### Brain-Behavior Correlations

We examined the relationship between neural stability (meanline) and synchronization consistency (standard deviation of the signed asynchronies). Figure 5 shows the simple correlations between participants’ EEG meanline values and the standard deviation of their signed asynchronies. Correlations were significant at all rates (SPR: *r*(22) = -0.43, *p* < 0.05; 15% Slower: *r*(22) = -0.65, *p* < 0.001; 30% Slower: *r*(22) = -0.43, *p* < 0.05). As can be seen in Figure 5, as meanline increased, the standard deviation of the signed asynchronies decreased.

**Figure 5.**
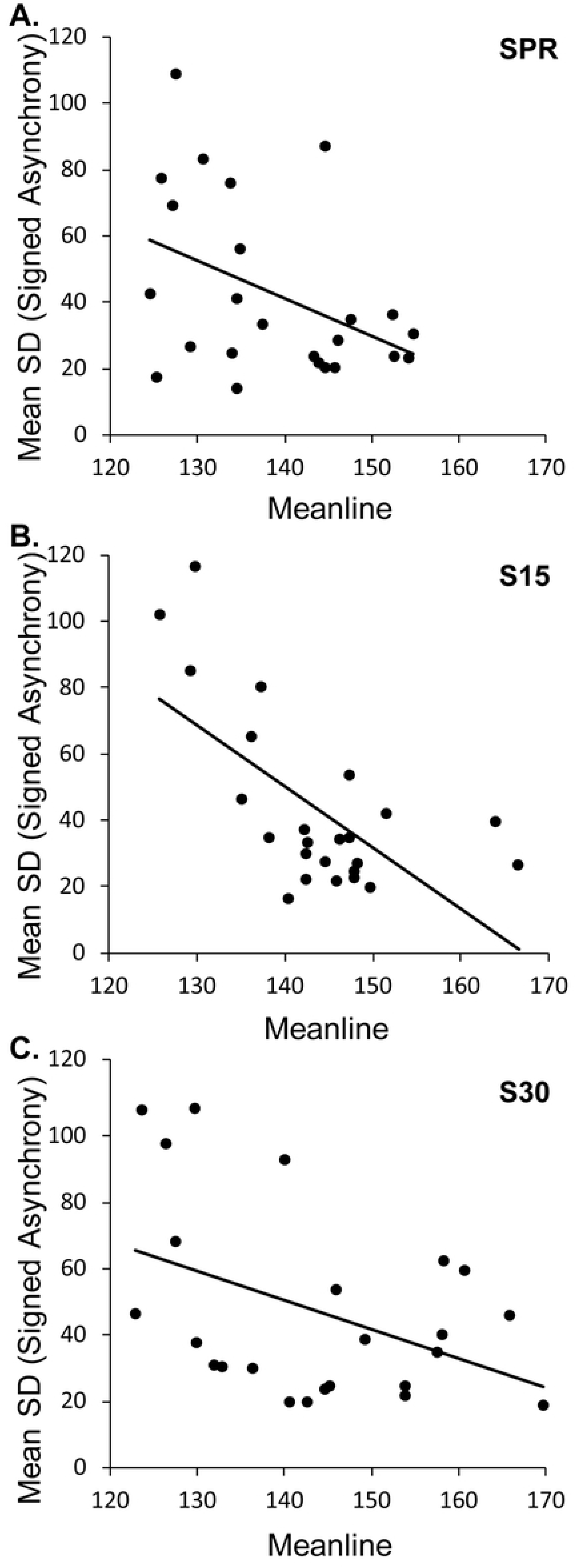
Correlations between EEG-based meanline (stability) values and standard deviation of participants’ asynchronies by rate condition. A) Spontaneous production rate; B) 15% Slower rate; and C) 30% Slower rate.

## Discussion

The current study explored the temporal dynamics of auditory-motor network activity during synchronization performance, and how these dynamics were influenced by musical training. First, synchronization accuracy decreased for both groups of participants as the tapping rate became slower than their spontaneous production rates. Second, musicians synchronized more accurately and more consistently than nonmusicians across rates, replicating previous research (e.g., [2, 19]). Musicians’ greater synchronization consistency is thought to reflect stronger auditory-motor integration [7]. This interpretation is consistent with our finding that musicians tended to show higher meanline (neural stability) than nonmusicians. Finally, brain-behavior comparisons showed significant negative correlations between meanline and the standard deviation of the signed asynchronies at all rates, indicating that neural stability increased as the synchronization consistency increased (ie, as the variability decreased).

Importantly, the meanline measure of neural stability in participants’ EEG signals was negatively correlated with the standard deviation of their signed asynchronies (indicating synchronization consistency) within each tapping rate. This is consistent with findings that suggest the strength of neural entrainment, measured via power in EEG activity at the stimulus frequency, is correlated with synchronization performance [44, 45]. Together, these findings suggest that neural stability may be a critical mechanism supporting auditory-motor synchronization across individuals and performance contexts. An advantage of recurrence quantification measures such as meanline and determinism is their sensitivity to nonlinear patterns of repetition typical of EEG signals as shown in Figure 1.

Neural predictability, measured by determinism of the EEG signal, increased for both participant groups as the rate became slower. Coordinating movements becomes more demanding at slower rates [46, 47]. The current study tailored the task demands to each individual by determining the synchronization rates based on participants’ comfortable tapping rates rather than fixing the rates across individuals. Task demands should be lowest at the individual’s SPR, representing their most comfortable rate, and increase as the rate becomes slower than their SPR. Previous findings suggest that increased task demands yield increased determinism in postural sway [48, 49]; our findings similarly showed increased deterministic structure of auditory-motor neural activity as the tapping rate became slower, reflecting increasing task demands. However, determinism (neural predictability) did not correlate with the mean signed asynchrony (synchronization accuracy). Future work could further investigate the relationship between neural predictability and behavioral outcomes.

The current findings suggest several future extensions. The RQA analysis was applied to recordings from a single electrode, based on previous findings that activity from this electrode best reflects auditory-motor integration [17]; contributions of other networks to the activity measured from this electrode are possible. Multivariate techniques such as multidimensional RQA can be applied to account for contributions from additional networks [50]. For example, Hall et al.[51] applied mdRQA to auditory-motor synchronization between individuals who phase-shifted their rhythms relative to each other to discover periods of behavioral stability. Source localization techniques could also be used to extract source activity from auditory and motor networks on which recurrence quantification analyses could be applied, and techniques such as magnetoencephalography could be used to obtain a better compromise between spatial and temporal resolution. Finally, extensions of this novel analysis to larger datasets may provide more discrimination between musicians’ and nonmusicians’ auditory-motor networks.

In sum, all participants showed decreased synchronization accuracy as the synchronization rate became slower, and musicians synchronized more consistently than nonmusicians across rates. Musicians tended to show greater neural stability than nonmusicians. Additionally, the predictability of auditory-motor neural activity increased as the rate became slower (as task demands increased), independent of musical training. Most importantly, for all participants, stability of the neural activity (measured with RQA) increased as consistency of behavioral synchronization increased. These findings support the interpretation that musical training leads to increased coactivation of auditory and motor brain regions, resulting in enhanced auditory-motor synchronization that is in turn associated with increased stability and predictability of temporal dynamics measured in electroencephalography. Future studies may extend the neural dynamics metrics of meanline and determinism to short-term (within-experiment) practice outcomes in a musician-nonmusician continuum.

## Declarations

The research was reviewed by the McGill University Research Ethics Board and the study was conducted in accordance with the ethical standards of the 1964 Declaration of Helsinki. The authors have no relevant financial or non-financial interests to disclose.

## Open Practices Statement

The datasets for the current study are available online in an open-source format.

## Acknowledgements

The research was funded by a Discovery Grant from the Natural Sciences and Engineering Research Council and by a Canada Research Chair to C. Palmer. We thank Sandra Kuliver for assistance. Correspondence can be addressed to C. Palmer, Dept of Psychology, McGill University, Montreal Canada H3A 1B1 or to.

## Notes

### Competing Interest Statement

The authors have declared no competing interest.

## References

[1] Mathias, B., Zamm, A., Gianferrara, P. G., Ross, B., & Palmer, C. Rhythm complexity modulates behavioral and neural dynamics during auditory-motor synchronization. J Cogn Neurosci. 2020; 32: 1864–1880.

[2] Scheurich, R., Zamm, A., & Palmer, C. Tapping into rate flexibility: Musical training facilitates synchronization around spontaneous production rates. Front Psychol. 2018; 9: 458.

[3] Tranchant, P., Scholler, E. & Palmer, C. Endogenous rhythms influence musicians’ and non-musicians’ interpersonal synchrony. Sci Rep. 2022; 12: 12973.

[4] Cameron, D. J., & Grahn, J. A. Enhanced timing abilities in percussionists generalize to rhythms without a musical beat. Front Hum Neurosci. 2014; 8: 1003.

[5] Krause, V., Pollok, B., & Schnitzler, A. Perception in action: The impact of sensory information on sensorimotor synchronization in musicians and nonmusicians. Acta Psychol (Amst). 2010; 133(1): 28–37.

[6] Zatorre, R. J., Chen, J. L., & Penhune, V. B. When the brain plays music: Auditory-motor interactions in music perception and production. Nat Rev Neurosci. 2007; 8(7): 547–558.

[7] Palomar-García, M-Á., Zatorre, R. J., Ventura-Campos, N., Bueichekú, E., & Ávila, C. Modulation of functional connectivity in auditory-motor networks in musicians compared with nonmusicians. Cereb Cortex. 2017; 27(5): 2768–2778.

[8] Bailey, J. A., Zatorre, R. J., & Penhune, V. B. Early musical training is linked to gray matter structure in the ventral premotor cortex and auditory-motor rhythm synchronization performance. J Cogn Neurosci. 2014;26: 755–767.

[9] Chen, J. L., Penhune, V. B., & Zatorre, R. J. The role of auditory and premotor cortex in sensorimotor transformations. Ann N Y Acad Sci. 2009; 1169(1): 15–34.

[10] Lahav, A., Saltzman, E., & Schlaug, G. Action representation of sound: Audiomotor recognition network while listening to newly acquired actions. J Neurosci. 2007; 27(2): 308–314.

[11] Bangert, M., & Altenmüller, E. O. Mapping perception to action in piano practice: A longitudinal DC-EEG study. BMC Neurosci. 2003; 4(1): 26.

[12] Mathias, B., Palmer, C., Perrin, F., & Tillmann, B. Sensorimotor learning enhances expectations during auditory perception. Cereb Cortex. 2015; 25(8): 2238–2254.

[13] Stober, S., Cameron, D. J., and Grahn, J. A. Using convolutional neural networks to recognize rhythm stimuli from electroencephalography recordings. Adv Neural Inf Process Syst. 2014; 27: 1449–1457.

[14] Marwan, N., & Webber, C. L. Mathematical and computational foundations of recurrence quantifications. In: Webber, C. Jr., Marwan, N., editors. Recurrence Quantification Analysis: Theory and Best Practices. Cham, Switzerland: Springer International Publishing; 2015. p 3–43.

[15] Acharya, U. R., Sree, S. V., Chattopadhyay, S., Yu, W., & Ang, P.C.A. Application of recurrence quantification analysis for the automated identification of epileptic EEG signals. Int J Neural Syst. 2011; 21: 199–211.

[16] Becker, K., Schneider, G., Eder, M., Ranft, A., Kochs, E. F., Zieglgänsberger, W., & Dodt, H. U. Anesthesia monitoring by recurrence quantification analysis of EEG data. PLoS One. 2010; 5(1): e8876.

[17] Scheurich, R., Demos, A. P., Zamm, A., Mathias, B., & Palmer, C. Capturing intra- and inter-brain dynamics with recurrence quantification analysis. In Proc Annu Conf Cogn Sci Soc. 2019a; 2748–2754.

[18] Zamm, A., Wang, Y., & Palmer, C. Musicians’ natural frequencies of performance display optimal temporal stability. J Biol Rhythms. 2019; 33: 432–440.

[19] Repp, B. H., & Doggett, R. Tapping to a very slow beat: A comparison of musicians and nonmusicians. Music Percept. 2007; 24(4): 367–376.

[20] Nozaradan, S., Zerouali, Y., Peretz, I., & Mouraux, A. Capturing with EEG the neural entrainment and coupling underlying sensorimotor synchronization to the beat. Cereb Cortex. 2015; 25(3): 736–747.

[21] Slater, J., Ashley, R., Tierney, A., & Kraus, N. Got rhythm? Better inhibitory control is linked with more consistent drumming and enhanced neural tracking of the musical beat in adult percussionists and nonpercussionists. J Cogn Neurosci. 2018; 30(1): 14–24.

[22] Oldfield, R. C. The assessment and analysis of handedness: The Edinburgh inventory. Neuropsychologia. 1971; 9(1): 97–113.

[23] Finney, S. A. FTAP: A Linux-based program for tapping and music experiments. Behav Res Methods Instrum Comput. 2001; 33(1): 65–72.

24. [24] Kothe, C. Lab streaming layer (LSL). 2014. Available from: https://github.com/sccn/labstreaminglayer.

[25] Zamm, A., Palmer, C., Bauer, A-K. R., Bleichner, M. G., Demos, A. P., & Debener, S. Synchronizing MIDI and wireless EEG measurements during natural piano performance. Brain Res. 2017; 1716: 27–38.

[26] Loehr, J. D., & Palmer, C. Temporal coordination between performing musicians. Q J Exp Psychol. 2011; 64(11): 2153–2167.

[27] Palmer, C. Sequence memory in music performance. Curr Dir Psychol Sci. 2005; 14: 247–250.

[28] Zamm, A., Pfordresher, P. Q., & Palmer, C. Temporal coordination in joint music performance: Effects of endogenous rhythms and auditory feedback. Exp Brain Res. 2015; 233(2): 607–615.

[29] Pecenka, N., & Keller, P. E. The role of temporal prediction abilities in interpersonal sensorimotor synchronization. Exp Brain Res. 2011; 211: 505–515.

[30] Scheurich, R., Pfordresher, P. Q., & Palmer, C. Musical training enhances temporal adaptation of auditory-motor synchronization. Exp Brain Res. 2019b; 238: 81–92.

[31] Fisher, N. I. Statistical analysis of circular data. Cambridge, UK: Cambridge University Press; 1993.

[32] Dalla Bella, S., & Sowiński, J. Uncovering beat deafness: Detecting rhythm disorders with synchronized finger tapping and perceptual timing tasks. J Vis Exp. 2015; 97:51761.

[33] Kirschner, S., & Tomasello, M. Joint drumming: Social context facilitates synchronization in preschool children. J Exp Child Psychol. 2009; 102(3): 299–314.

[34] Sowiński, J., & Dalla Bella, S. Poor synchronization to the beat may result from deficient auditory-motor mapping. Neuropsychologia. 2013; 51(10): 1952–1963.

[35] Delorme, A., & Makeig, S. EEGLAB: An open source toolbox for analysis of single-trial EEG dynamics including independent component analysis. J Neurosci Methods. 2004; 134(1): 9–21.

[36] Marwan, N., Romano, M. C., Thiel, M., & Kurths, J. Recurrence plots for the analysis of complex systems. Phys Rep. 2007; 438(5-6): 237–329.

[37] López Pérez, D, Bokde, ALW, & Kerskens, CM. Complexity analysis of heartbeat-related signals in brain MRI time series as a potential biomarker for ageing and cognitive performance. Eur Phys J Spec Top. 2023; 232: 123–133.

[38] Demos, A.P., Chaffin, R., & Logan, T. Musicians’ body sway embodies musical structure and expression: A recurrence-based approach. Musicae Scientiae. 2017; 22: 244–263.

[39] Laudańska, D., Radkowska, A., Babis, K., Malinowska-Korczak, A., Wallot, S., & Tomalski, P. Changes in the complexity of limb movements during the first year of life across different tasks. Entropy. 2022; 24: 552. 10.3390/e24040552

[40] Rosen, P. J., Epstein, J. N., & Van Orden, G. I know it when I quantify it: Ecological momentary assessment and recurrence quantification analysis of emotion dysregulation in children with ADHD. Atten Defic Hyperact Disord. 2013; 5(3): 283–294.

[41] Holm, S. A simple sequentially rejective multiple test procedure. Scan J Stat. 1979; 6: 65–70. [44] Bouvet, C. J., Bardy, B. G., Keller, P. E., Dalla Bella, S., Nozaradan, S., & Varlet, M. Accent-induced modulation of neural and movement patterns during spontaneous synchronization to auditory rhythms. J Cogn Neurosci. 2020; 32(12): 1–12.

[42] Levene, H. Robust tests for equality of variances. In: Olkin, I. Ghurye, S. G. Hoeffding W. Madow W. G. Mann H. B., editors. Contributions to probability and statistics: Essays in honor of Harold Hotelling. Stanford: Stanford University Press; 1960. pp. 278–292.

[43] Mair, P., & Wilcox, R. Robust statistical methods in R using the WRS2 package. Behav Res Methods. 2020; 52: 464–488.

[44] Bouvet, C. J., Bardy, B. G., Keller, P. E., Dalla Bella, S., Nozaradan, S., & Varlet, M. Accent-induced modulation of neural and movement patterns during spontaneous synchronization to auditory rhythms. J Cogn Neurosci. 2020; 32(12): 1–12.

[45] Nozaradan, S., Peretz, I., & Keller, P. E. Individual differences in rhythmic cortical entrainment correlate with predictive behavior in sensorimotor synchronization. Sci Rep. 2016; 6: 20612.

[46] Fujiyama, H., Hinder, M. R., Garry, M. I., & Summers, J. J. Slow and steady is not as easy as it sounds: Interlimb coordination at slow speed is associated with elevated attentional demand especially in older adults. Exp Brain Res. 2013; 227(2): 289–300.

[47] McPherson, T., Berger, D., Alagapan, S., & Fröhlich, F. Intrinsic rhythmicity predicts synchronization-continuation entrainment performance. Sci Rep. 2018; 11782.

[48] Balasubramaniam, R., Riley, M. A., & Turvey, M. T. Specificity of postural sway to the demands of a precision task. Gait Posture. 2000; 11(1), 12–24.

[49] Mazaheri, M., Salavati, M., Negahban, H., Sanjari, M. A., & Parnianpour, M. Postural sway in low back pain: Effects of dual tasks. Gait Posture. 2010; 31(1): 116–121.

[50] Wallot, S., Roepstorff, A., & Mønster, D. Multidimensional Recurrence Quantification Analysis (MdRQA) for the analysis of multidimensional time-series: A software implementation in MATLAB and its application to group-level data in joint action. Front Psychol. 2016; 7: 1835.

[51] Hall C., Kim, J.C., & Paxton, A. Multidimensional recurrence quantification analysis of human-metronome phasing. PLoS One. 2023; 18: e0279987.

